# Antiproliferative Effects of Novel Copper (II) Complexes on Lung Cancer Cell Line

**DOI:** 10.1101/2022.08.12.503805

**Authors:** Muhammed Fawaz Abdullah, Nilufer Cinkilic, Ozgur Vatan, Duygu Inci, Rahmiye Aydin

## Abstract

Copper is an essential metalloelement that plays key fundamental roles in both health and pathology, and is increasingly been implicated in molecular pathogenesis of many cancer types. It has shown promise as a replacement to cisplatin in coordination complexes presently in mainstream chemotherapeutic practices.

In this study, two newly synthesized water-soluble ternary copper (II) mixed ligand complexes; complex 1 - (Cu(4-mphen)(tyr)(H2O)]NO3·2H2O)(**C.1**) and complex 2 - (Cu(5-mphen)(tyr)(H2O)]NO3·2H2O (**C.2**) where (4-m= 4-methyl; 5-m = 5-methyl; phen-1, 10 = phenanthroline; tyr = tyrosine)), were investigated on adenocarcinomic human alveolar basal epithelial cell, A549 and non-cancerous human bronchial epithelial cell, BEAS-2B for their antiproliferative effects using the XTT assay (cytotoxicity), Comet assay (genotoxicity) and DCFH-DA assay (intracellular ROS) tests.

C.1 was significantly more cytotoxic in A549 than C.2. Data from the Comet and ROS assay tests support each other. C.2 caused more copper-induced DNA damage, possibly through significant induction of ROS-mediated oxidative damage in the cancer cell, but a minimal insignificant ROS rise in normal cells. These results can only be preliminary and further studies are required to better understand the cellular effects and functional interactions of these agents, for an efficient therapeutic design and application.

## INTRODUCTION

Copper complexes are increasingly gaining recognition as metal-based drug candidates in cancer therapeutics due to their broader spectra of activities (bioavailability, wide structural variability, biologically accessible redox abilities and lower toxicity (1–3), providing a possible means of escaping the clinical issues encountered by approved platinum-based drugs. It has become quite imperative in anticancer drug development to consider compounds with increased efficacy and potentially reduced harmful side effects. Collateral suicidal damage of anticancer drugs to normal cells is a primary cause of their side effects. Cisplatin though relatively successful, is limited due to its toxic side effects and development of cellular resistance (38). Copper as well as other endogenous metal ions compared to non-endogenous metals (cisplatin, platinum-based anticancer agents, etc.) tend to be less toxic to normal cells. These motivated the search for new metal-based anti-cancer agents with selectivity activity against cancer cells.

Copper, described as a modern bio-element (4–6) is a well-known redox metal with critical cellular functions (7, 8). These critical cellular roles have identified an abnormal accumulation of copper in cancer cells, a distinctive feature in transformed and healthy cells, and which can be targets for new chemotherapeutic agents (9). Designed compounds targeting these redox process and thus modulate intracellular reactive oxygen species (ROS) levels can be beneficial.

To enhance biological activity in compound synthesis, Cu (II) complexes are commonly bound to biologically active ligands such as complexes of thiosemicarbazones (10) to Cu (II). Effects synthetic copper (II) complexes and potential anticancer agents as pharmacological agents have been reported (11; 12). Very recently, some mixed ligand copper (II) complexes, which strongly bind and cleave DNA, exhibit significant anti-cancer activities and have been shown to regulate apoptosis (13). Copper complexes are available for a broader spectrum of antitumor activity (14). Their distinct mechanism of pharmacological actions different from platinum-based agents (cisplatin), Copper complexes have shown potential to replace platinum-based agents (cisplatin) in coordination complexes presently in mainstream chemotherapeutic practices.

Phenanthroline (phen) ligand and its derivatives are famous for their metal-chelating properties (15, 16) and have widely used in biological probes development and numerous analytical reagents (17) since the discovery of its nuclease activity (18) or ts DNA intercalation abilities. Tyrosine, like other redox active aromatic amino acids are significantly used in drug synthesis, with their aromatic interaction having major function in biomolecular recognition processes in metalloenzyme-catalysed substrate oxidations (19–21)

Current interest in copper complexes is based on their potential use as antimicrobial, antiviral, anti-inflammatory, anti-tumor agents, enzyme inhibitors or chemical nucleases. It has been shown that copper accumulates in the tumor cell due to the selective permeability of the cancer cell membrane. Numerous copper complexes have been studied for anti-cancer activity, and some have been found to be effective both in vivo (in mouse tumor models) and in vitro (on cancer cell lines in culture). In addition, copper (II) complex appears to be a promising candidate for anticancer therapy, at which point there are many studies describing the synthesis and cytotoxic activity of copper (II) complexes (22, 23).

Numerous studies have described copper (II) complexes containing phen or substituted phen derivatives and L-amino acids to be potent cytotoxic agents, possessing sub-micromolar range IC_50_ values. For instance, [Cu(phen)_2_](NO_3_)_2_ was shown to display higher cytotoxic effect at 24 h (IC_50_ - 1.1lM) in PC3 (prostate cancer cell line) than at 96 h (IC_50_ - 15.2lM), while in PNT1A (noncancerous prostate cell line) and SK-OV-3 (ovaries cancer cell line), IC_50_ were 1.8 and 0.6 lM, respectively at the 96 h (24). Similarly, Pivetta and coworkers synthesized two binary copper (II) complexes (Cu(phen)(OH_2_)_2_(ClO_4_)_2_ and [Cu(phen)_2_(OH_2_)](ClO_4_)_2_) and studied its cytotoxic activity in four (4) human cancer-derived cell lines (CCRF-CEM acute T-lymphoblastic leukemia, CCRF-SB acute B-lymphoblastic leukemia, K-MES-1 lung squamous carcinoma and DU-145 prostate carcinoma), with IC_50_ in the 1-3l M range. Due to its two phen units, [Cu(phen)_2_(OH_2_)](ClO_4_)_2_ was more cytotoxic than Cu(phen)(OH_2_)_2_(ClO_4_)_2_ (25).

In another study, the anti-cancer effects of copper (II) mixed complexes with [Cu(L-tyr)(diimine)]ClO_4_, 1,10-phenanthroline, dimethyl-1,10-phenanthroline and dipyridoquinoxaline ligands were investigated in NCI-H460 lung cancer cell lines (26).

Copper complexes with cisplatin (covalent DNA binding) are of interest based on different modes of action. It includes DNA interference, mitochondrial toxicity and ROS production (27). Cu (II) complexes are considered the most promising among cisplatin as anticancer agents. However, little understanding of the molecular basis of its mechanism of action of Copper (II) and thus copper (II) mixed ligands as a pharmacological agent in biological systems has been documented. This study therefore seeks to understand the molecular bases of the biomolecular interactions of copper (II) complexes with DNA, protein and other biological entities and thus chemical structure-biological activity relationship. Deciphering the modes of metal complexes-DNA binding is essential to finding the fundamental principles governing biomolecular recognition processes (affinity and specificity of chemical complexes) of these functional agents.

## MATERIALS AND METHODS

Different effects of two novel water-soluble ternary copper (II) complexes with mixed ligands: **Complex 1**: [Cu(4-mphen)(tyr)(H_2_O)]ClO_4_ (**C.1)** and **Complex 2**: [Cu_2_(5-mphen)_2_(tyr)2(H2O)_2_]·(ClO_4_)_2_·3H_2_O (**C.2)** in A549 (adenocarcinomic human alveolar basal epithelial cell) and BEAS-2B (immortalized non-cancerous human bronchial epithelial cell line) were evaluated using XTT assay (cytotoxic effect), alkaline comet test (genotoxic effects) and DFCD-DA test (intracellular reactive oxygen species (ROS)). (m – methyl; phen - 1,10-phenanthroline and tyr - tyrosine).

Cells were cultured in RPMI-1640 medium supplemented with 15% fetal calf serum (FCS), penicillin (100 IU/mL) and streptomycin (100 lg/ mL), 10 mM L-glutamine, 10 mM non-essential amino acids and sodium pyruvate, and maintained at 37 °C in a humidified atmosphere containing 5% CO_2_. A549 and BEAS-2B cell lines were grown in 75 cm^2^ flasks. The cell lines were further sub-cultured once a week. Grown cells were harvested and counted.

### Cytotoxic effects (XTT assay)

The cytotoxic effects of the two copper (II) complexes were evaluated against A549 and BEAS-2B cell lines was measured using the XTT cell viability kit, with cisplatin as the reference metallodrug (control). This assays the reduction of XTT (tetrazolium salt) to orange formazan colour by metabolic active cells and correlates it to cell viability.

Cells grown in 75 cm^2^ flasks and sub-cultured once weekly, were harvested and counted. A549 and BEAS-2B were respectively seeded at 5×10^3^ cells/well in sterile 96-well flat-bottomed plates in triplicate, including blanks and incubated overnight at 37 °C in a humidified atmosphere containing 5% CO_2_. The blanks were filled with the complete medium minus cells. The copper (II) complexes were dissolved in sterile distilled water. Cells were then treated with various concentrations (0.1 - 20 μM) of copper (II) complexes for 24 h. Cisplatin was also evaluated as a control, under the same experimental conditions.

After the 24 h treatment, wells were washed with PBS and fresh medium (100 μL) added. Per the manufacturer’s instructions (Biological Industries), activated XTT solution (50 μL) was added to each well and the plate incubated for another 3 hours in the CO_2_ incubator at 37 °C. Colour change absorbance was read at a wavelength of 450 nm with a microplate reader, and cell proliferation curve drawn to further calculate the half-maximal inhibitory concentration (IC_50_).

After blank subtraction, the percent growth inhibition of cells was calculated as follows:

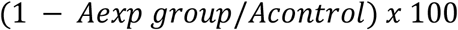

### Genotoxic damage (Alkaline Comet assay)

The alkaline comet assay detected the single strand DNA breaks induced by our copper complexes. Cells were treated at 310.2 K for 24 hours with IC_12.5_, IC_25_, IC_50_ and IC_75_ doses of copper complexes, in duplicates. Cells were harvested after the 24 hours treatment, and suspended in 0.75% low melting agarose dissolved in PBS. The cells-agarose mixture was then spread onto 0.8% normal melting point agarose-pre-coated slides and kept overnight in lysis solution (2.5 M NaCl, 0.1 M EDTA, 10 mM Tris base, pH 10, 1% Triton X, 10% DMSO) at 4 °C. Slides were further subjected to DNA unwinding for 20 min in the alkaline buffer (300 mM NaOH, 1 mM EDTA, pH > 13). Electrophoresis was conducted for 20 min at 25 V (300 mA) in alkaline buffer. Slides were immersed in neutralization buffer (pH = 7) for 5 min following electrophoresis and then dehydrated in 70% ethanol. The slides stained with ethidium bromide, were analyzed under Nikon epi-fluorescence microscope (Eclipse 80i) equipped with a digital camera (Kameram 21, Istanbul, Turkey). Quantitative assessment of DNA damage was performed with Comet software (Arganit Microsystems Comet Assay, Istanbul, Turkey) in 200 randomly selected nuclei per treatment. % Tail DNA (TDNA), tail length (TL) and Olive tail moment (OTM) values of each comet were measured.

### Detection of reactive oxygen species (ROS) induction

Abilities of the two copper complexes to induce intracellular ROS was assessed using the 2’,7’-dichlorodihydrofluorescein diacetate (DCFH-DA) assay. This dye itself influorescent, upon diffusion into cells and subsequent hydrolysis by cellular esterase transform into DCFH, and rapidly oxidized to the highly fluorescent 2’,7’-dichlorodihydrofluorescein (DCF) in the presence of intracellular ROS.

Cells grown in black 96-well cell culture plates in duplicates, were pre-incubated with DCFH-DA for 2 hrs and washed twice with DPBS afterwards. DCFH-DA-loaded cells were treated with IC_12.5_, IC_25_, IC_50_ and IC_75_ doses of copper complexes for two hours, with hydrogen peroxide (H_2_O_2_) as the positive control. A 1:10 dilution series of DCF standards ranging in concentration from 0 μM to 10 μM is prepared by tube dilution method of the 1mM DCF stock solution with the medium. 10 μl of the DCF stock is taken and added to the 1st standard containing 990 μl of RPMI medium, and then 100 μl is taken from standard 1 to the other standards and diluted. The DCF fluorescence intensity of each well was measured with an excitation wavelength of 480 nm and an emission wavelength of 530 nm using a standard fluorescence plate reader (Thermoscientific Fluoroskan Ascent FL). The ROS contents were determined using Relative Fluorescent Units (RFU). The experiment was carried out in duplicate. Statistical analyzes were performed with the Mann-Whitney U test using the SPSS 24 package program.

## RESULTS AND DISCUSSION

The fundamental aim of any anticancer drug development are essentially to increase efficacy of a potential drug agent while reducing its harmful side effects. Cytotoxic effects of various anticancer drugs to normal cells underlie their major side effects. These toxic effects and emergence of cellular resistance explains cisplatin’s limited use, necessitating the need for new metal-containing anti-cancer agents with selective properties against cancer cells. Our work thus studied the antiproliferative effects of two newly synthesized copper (II) mixed ligand complexes**: C.1**and **C. 2** in A549 and BEAS-2B cell lines by measuring cytotoxicity, genotoxicity and ROS-mediated oxidative damage.

Previous studies have described copper (II) complexes containing phen or substituted phen derivatives and L-amino acids to be potent cytotoxic agents, possessing sub-micromolar range IC_50_ values. For instance, [Cu(phen)_2_](NO_3_)_2_ was shown to display higher cytotoxic effect at 24 h (IC_50_ - 1.1lM) in PC3 (prostate cancer cell line) than at 96 h (IC_50_ - 15.2lM), while in PNT1A (noncancerous prostate cell line) and SK-OV-3 (ovaries cancer cell line), IC_50_ were 1.8 and 0.6 lM, respectively at the 96 h (24). Similarly, Pivetta and coworkers synthesized two binary copper (II) complexes (Cu(phen)(OH_2_)_2_(ClO_4_)_2_ and [Cu(phen)_2_(OH_2_)](ClO_4_)_2_) and studied its cytotoxic activity in four (4) human cancer-derived cell lines (CCRF-CEM acute T-lymphoblastic leukemia, CCRF-SB acute B-lymphoblastic leukemia, K-MES-1 lung squamous carcinoma and DU-145 prostate carcinoma), with IC_50_ in the 1-3l M range. Due to its two phen units, [Cu(phen)_2_(OH_2_)](ClO_4_)_2_ was more cytotoxic than Cu(phen)(OH_2_)_2_(ClO_4_)_2_ (25).

In determining the possible cytotoxic effect of our newly synthesized compounds against cellular metabolism (cell viability), in vitro XTT tests were performed on A549 (lung tumour cell line) and BEAS-2B (normal cell). Cisplatin was also similarly evaluated for comparative analysis. Their IC_50_ values were determined relative the in vitro cytotoxic effects. **C.1** recorded IC_50_ values of 1.727 and 2.880 in A549 and BEAS-2B respectively, while **C.2** had IC_50_ values of 2.812 in A549 and 2,828 in BEAS-2B (**Tab. 1**).

**Table 1:** The cytotoxic activities of the complexes

| Compounds | Cell lines ($IC_{50}$ (μM)) | |
| --- | --- | --- |
|  | A549 | BEAS-2B |
| **C.1** | 1,727 ± 0,190* | 2,880 ± 0,299* |
| **C. 2** | 2,812 ± 0,24 | 2,828 ± 0,253 |
| Cisplatin | 22.2 ± 0.4 | 18.9 ± 0.9 |

**C.1** exhibited higher cytotoxic potential against the lung cancer cell at much lower IC_50_ value, indicating greater ability to eliminating the cancer cells even at low concentrations (**Fig. 2**). Inci and colleagues previously showed cisplatin to have IC_50_ values of 22.2 and 18.9 in A549 and Beas-2B cell lines respectively (28). Interestingly, the two copper complexes have lower IC_50_ values than cisplatin for the same cell line and thus potential to act as metal-based anticancer drugs.

**Fig. 2:**
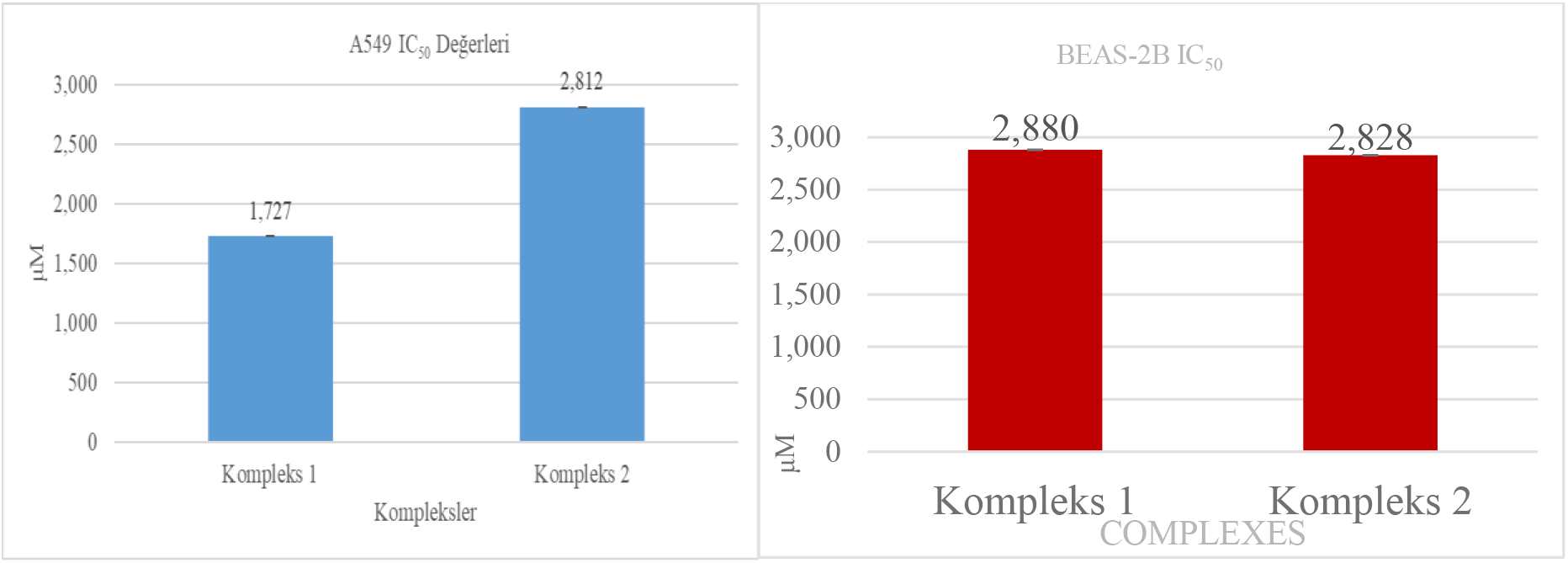
IC_50_ values (μM) of **C.1** (kompleks 1) & **C.2** (kompleks 2) as determined in A549 and BEAS-2B

Recently, Fei and colleagues investigated the anti-cancer effects of Copper (II) N-(pyridin-2-ylmethylene) dehydroabietylamine complexes in HeLa, SiHa, HepG-2 and A431 cancer cell lines, reported the complex has a very high cytotoxic effect compared to cisplatin (29). They particularly reported higher cytotoxicity in HeLa cells via apoptotic mechanism. In another study, the anti-cancer effects of copper (II) mixed complexes with [Cu(L-tyr)(diimine)]ClO_4_, 1,10-phenanthroline, dimethyl-1,10-phenanthroline and dipyridoquinoxaline ligands were investigated in NCI-H460 lung cancer cell lines (26). IC_50_ values of all the three complexes were determined between the range of 900-4900 nM. These values are in agreement with our IC_50_ findings.

Genetic damages induced by our copper (II) mixed ligand complexes were examined by alkaline comet assay. A549 and BEAS-2B cell lines treated with various concentrations (IC_12.5_, IC_25_, IC_50_ and IC_75_) of **C.1** and **C.2** for 24 hours. The three (3) important parameters, generally accepted for assessment of comet test findings: Tail length, Tail %DNA and Olive Tail Moment (OTM) were evaluated. Dose-dependent DNA damage was observed after cellular exposure to both **C.1** and **C.2**. Evident from the obtained data on the bases of DNA tail length (distance from DNA head to DNA tail), OTM and tail % DNA induced by the two complexes, significant DNA damage was observed in both cell lines even at the lowest dose. However, it was statistically low and insignificant in BEAS-2B cells. The rate of DNA damage especially in A549, also increases with increasing concentration (**Fig. 3**).

**Fig. 3:**
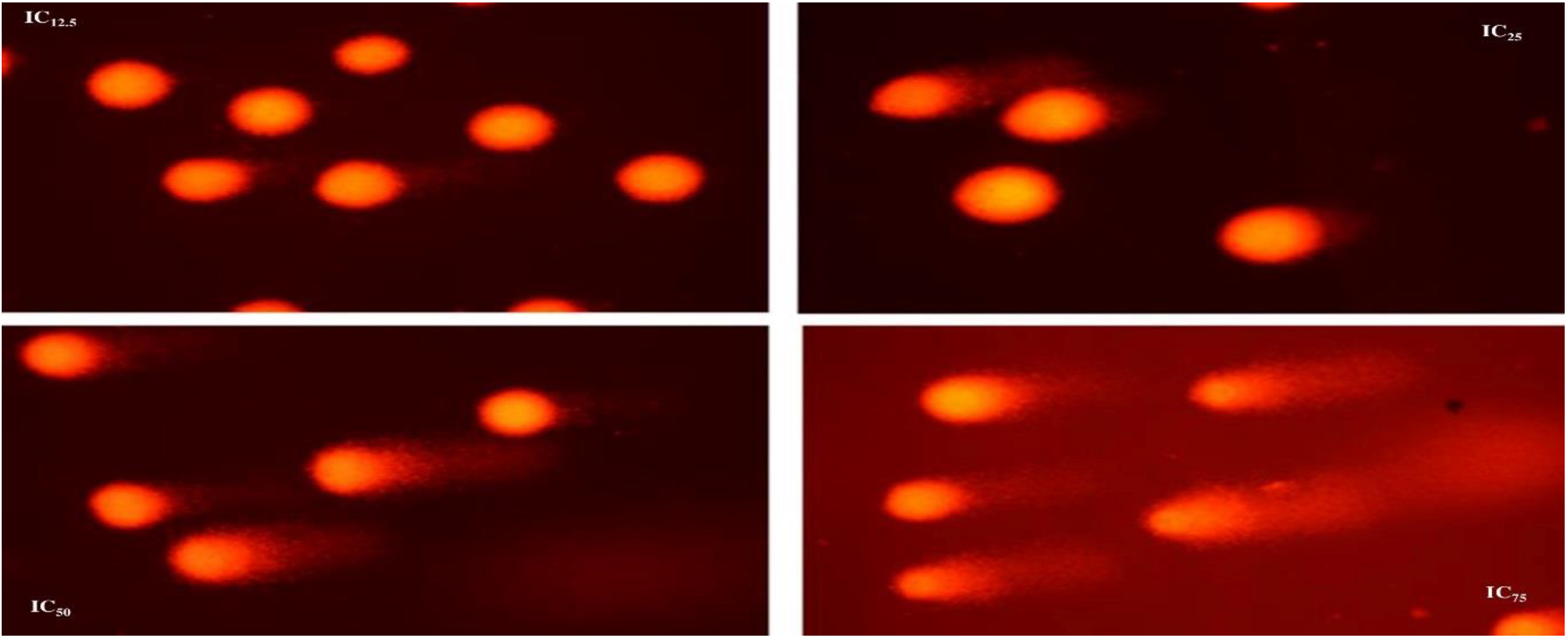
Photographic image of comet assay under fluorescence microscopy (X20) C.2-induced DNA damage in A549 cells treated with IC_12.5_, IC_25_, IC_50_, IC_75_ doses of the complex.

Assessment of our comet test results showed that **C.2** had more genotoxic effect in A549 and caused greater DNA damage statistically than **C.1 (Fig. 4)**. This is quite interesting on the basis of the higher IC_50_ value of **C.2.** Both complexes induced less genotoxic damage in BEAS-2B as anticipated. Our planar copper (II) mixed ligand complexes in quaternary structure, can interact with cells via DNA intercalation forming single and double strand DNA breaks (28). Our findings are in support of such interactions. These compounds can additionally increase the level of intercellular ROS (30). Oxidative damage is generally accepted to play a role in carcinogenesis. On the other hand, some anti-cancer drugs especially alkylating agents, are known to employ oxidative stress in their mechanism of action. Some studies have shown cancerous cells to be in increased oxidative state, and therefore very vulnerable to increased ROS formation (31). For example, the pro-oxidant effect of cisplatin includes inhibition of thioredoxin reductase and increased ROS production in mitochondria (32). The likely cause of the excessive production of ROS induced by cisplatin is the degradation of mitochondrial DNA (mtDNA), causing an imbalance in the expression of apolipoprotein in the mitochondrial respiratory chain and leading to leakage of superoxide (O_2_^•-^).

**Fig. 4:**
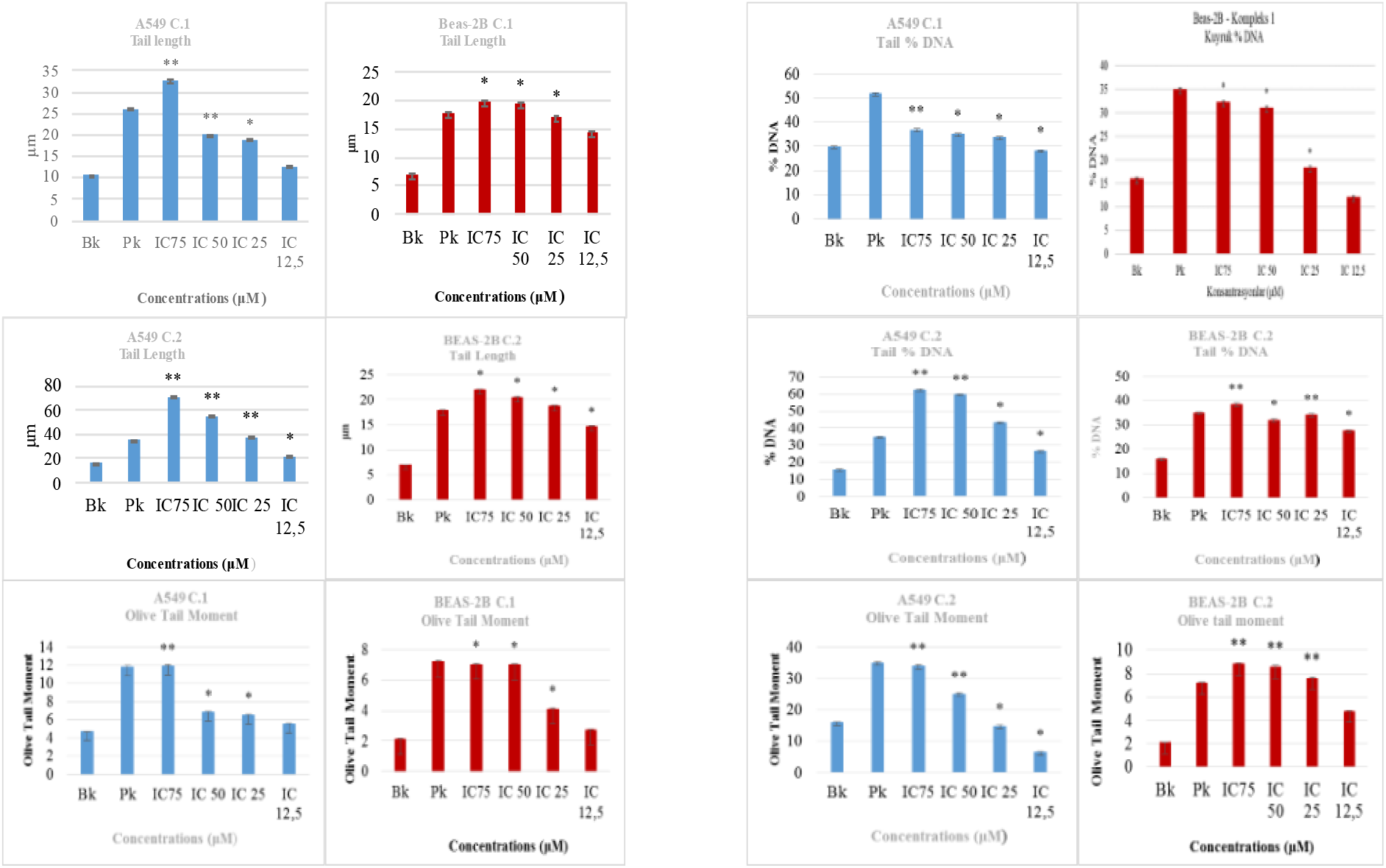
Comet assay results of A549 and BEAS-2B cell lines exposed to **C.1** and **C.2** (* p < 0.05, **p < 0.001)

Due to the high copper level and high oxidative stress in cancer cells, it has been reported that compounds that target the redox process and thus modulate the intracellular ROS level may be beneficial. Copper complexes are well known for their redox property and ROS production. However, most studies only involve cell-free systems (36). To examine the induction of ROS production by **C.1** and **C.2** in A549 and Beas-2B cell lines, both cell lines were treated at various concentrations (IC_12.5_, IC_25_, IC_50_ and IC_75_). **C. 2** was generally found to cause higher significant ROS production than **C.1**. In addition, the increase in ROS values in A549 (**Fig. 5** and **Fig. 6**) was observed at higher levels than in Beas-2B. This is a typical feature expected of any anti-cancer agent. ROS production of both complexes in Beas-2B were particularly observed to be almost equal. These findings showed that the complexes have the ability to increase intracellular and intercellular oxidative damage but less oxidative damage in healthy cells.

**Fig. 5:**
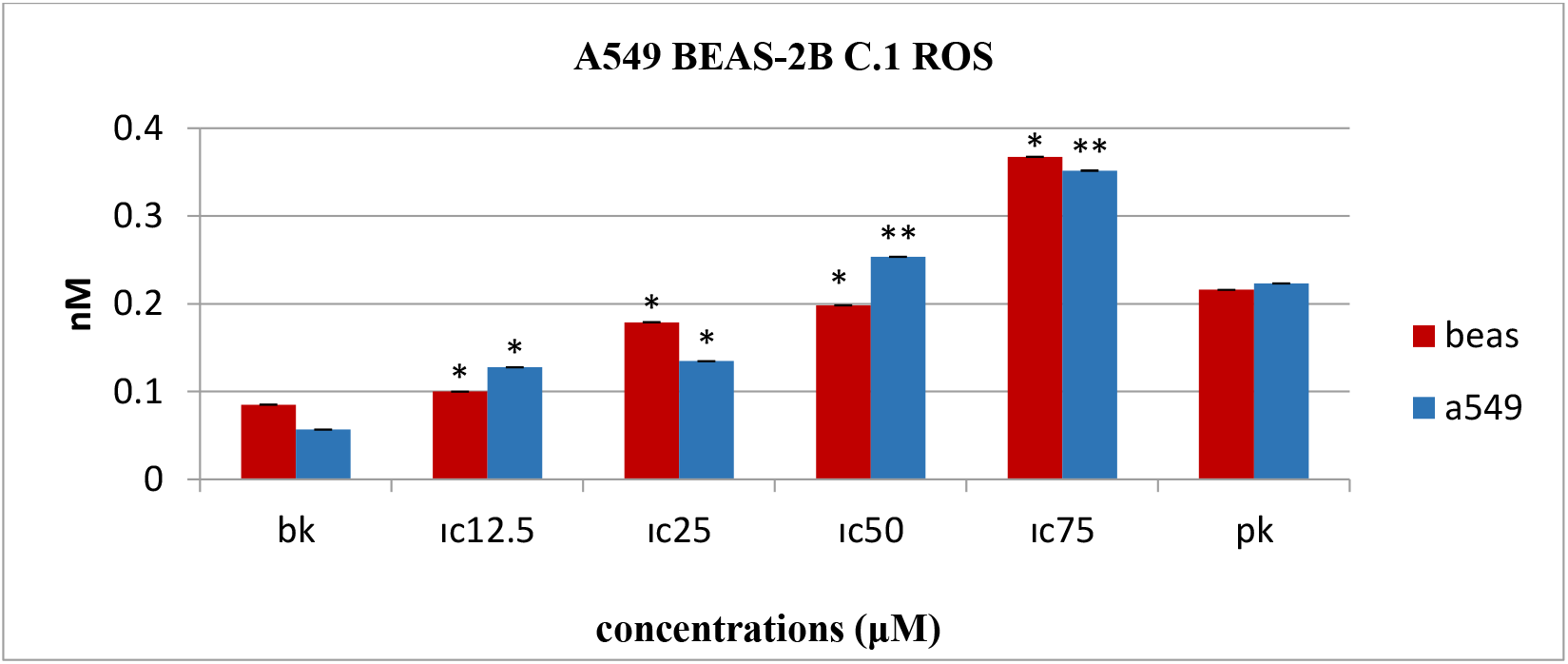
Effect of **C.1** on ROS generation in BEAS-2B ve A549 cell lines (* p < 0.05, **p < 0.001)

**Fig. 6:**
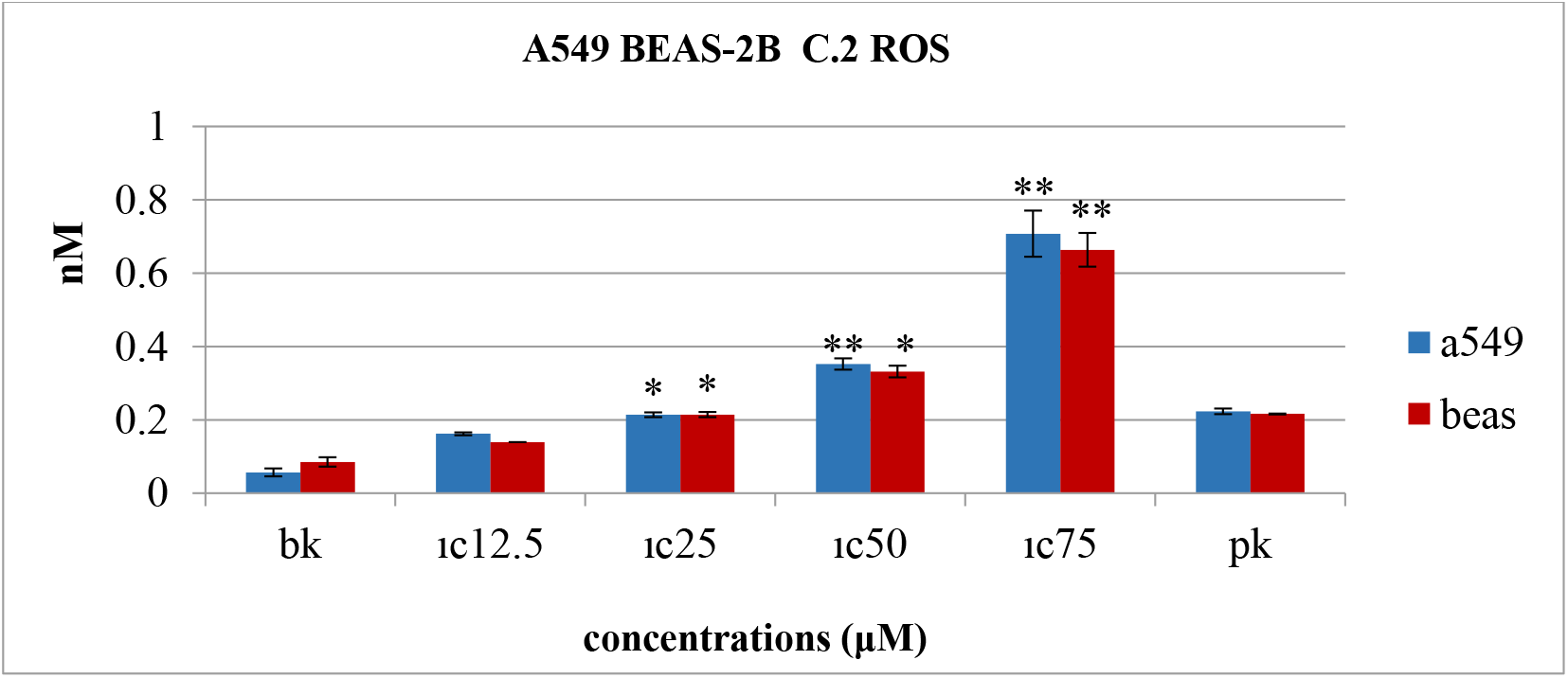
Effect of **C.2** on ROS generated A549 and BEAS-2B cell lines (* p < 0.05, **p < 0.001)

The type of radical produced, the region where the radical is located, and the local concentration are important in the increase of ROS levels in cancer. The observed high level of ROS in A549 supports the evidence for abnormally elevated ROS levels commonly observed in various cancer cells. In many types of cancer, ROS-sensitive signaling pathways involved in cell growth/proliferation, differentiation, protein synthesis, glucose metabolism, cell survival and inflammation are constantly increased. Reactive oxygen species, especially hydrogen peroxide, can act as second messengers in cellular signaling. H_2_O_2_ regulates protein activity by reversible oxidation of the target, such as protein tyrosine phosphatases, protein tyrosine kinases, receptor tyrosine kinases, and transcription factors (8, 33).

Increased metabolic activity of high ROS level in cancer cells causes mitochondrial dysfunction, peroxisome activity, increased cellular receptor signaling, oncogenic and thymidine phosphorylase activity. The tumor cell shields itself from ROS by increasing its antioxidant protein expression levels to ensure its safety. Thus, it is thought that a delicate balance of intracellular ROS levels is necessary for the function of the cancer cell. For example: cancer cell can adapt to elevated ROS level by increasing intracellular antioxidants such as glutathione (GSH) and heme oxygenase-1 (HO-1) (31, 34, 35).

## CONCLUSIONS

In this study, some antiproliferative (anticancer) effects of the two newly synthesized copper (II) mixed ligand complexes; **Complex 1**: [Cu(4-mphen)(tyr)(H_2_O)]ClO_4_(**C.1)** and **Complex 2**: [Cu_2_(5-mphen)_2_(tyr)2(H_2_O)_2_]·(ClO_4_)_2_·3H_2_O(**C.2)** in human lung cancer (A549) and normal human lung epithelial cell lines (BEAS-2B) were evaluated. These two complexes were determined to cause varying degrees of cytotoxic, genotoxic and oxidative damage in the cancer cell line and were more potent than cisplatin at most experimental concentrations. These varying degrees of effects can be said to be a product of their mechanism of action and chemical structure, which determine the nature of their cellular biomolecular interactions. By virtue of the cumulative effect of chemical structure and biomolecular interactions, **C.1** had more cytotoxic effect selectively in lung cancer cells but lesser ability to induce genetic damage. The exact opposite was the case in **C.2**. Copper-induced damage and ROS-mediated oxidative damage augmented each other.

These distinctively different behaviours in their pharmacological action indicate the need for further studies to elucidate their selectivity, target, mode of action (biomolecular interactions), its applicability in other cell lines and in vivo effects. This will aid in our quest to find alternative drug candidates that can overcome disease (cancer) resistance.

## ACKNOWLEDGEMENT

We wish to acknowledge the financial support of **Bursa Uludag University Research Fund** for these works (**Project Numbers OUAP (F)-2015/14**).

## Supplementary list

**Figures 4:**
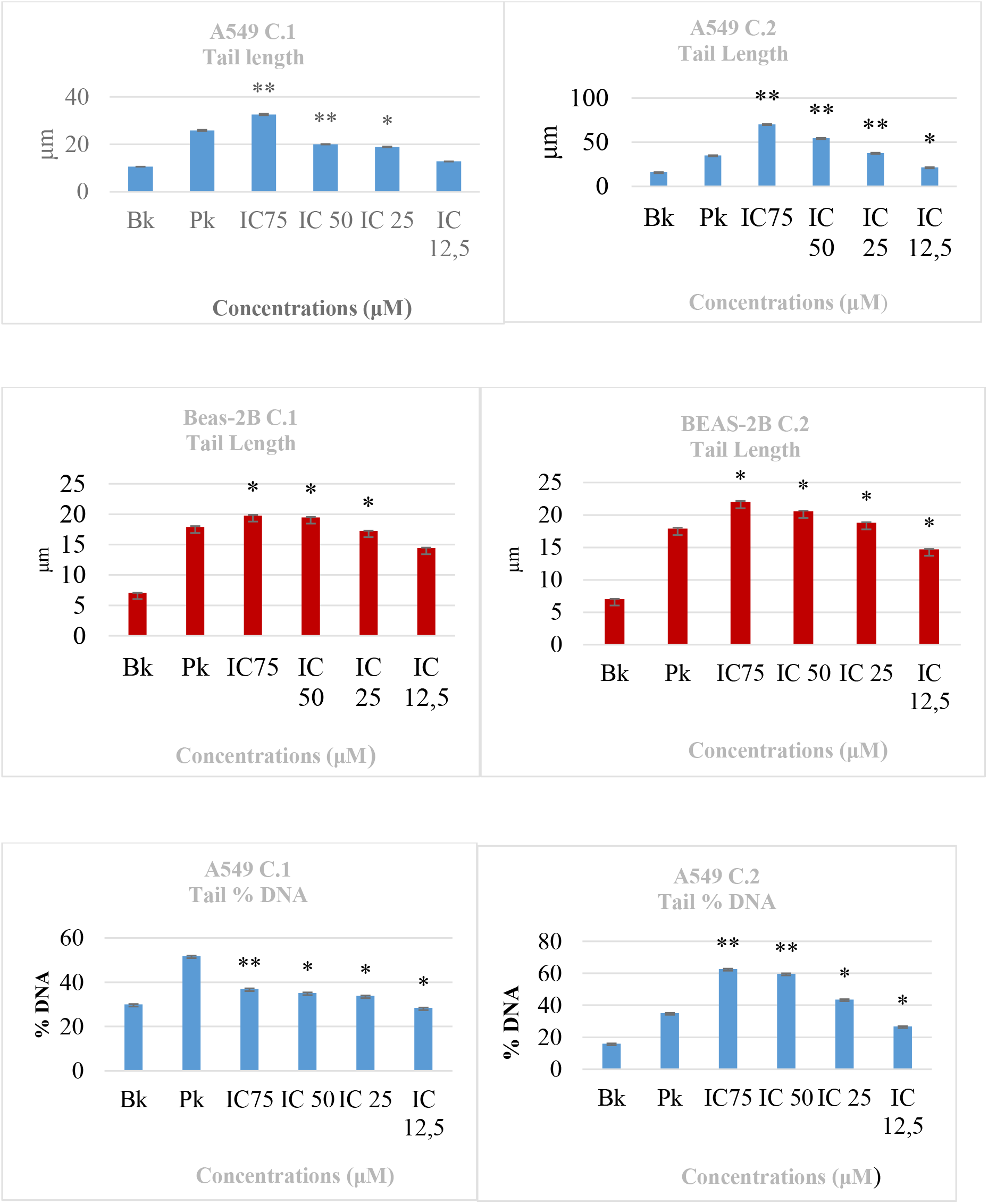
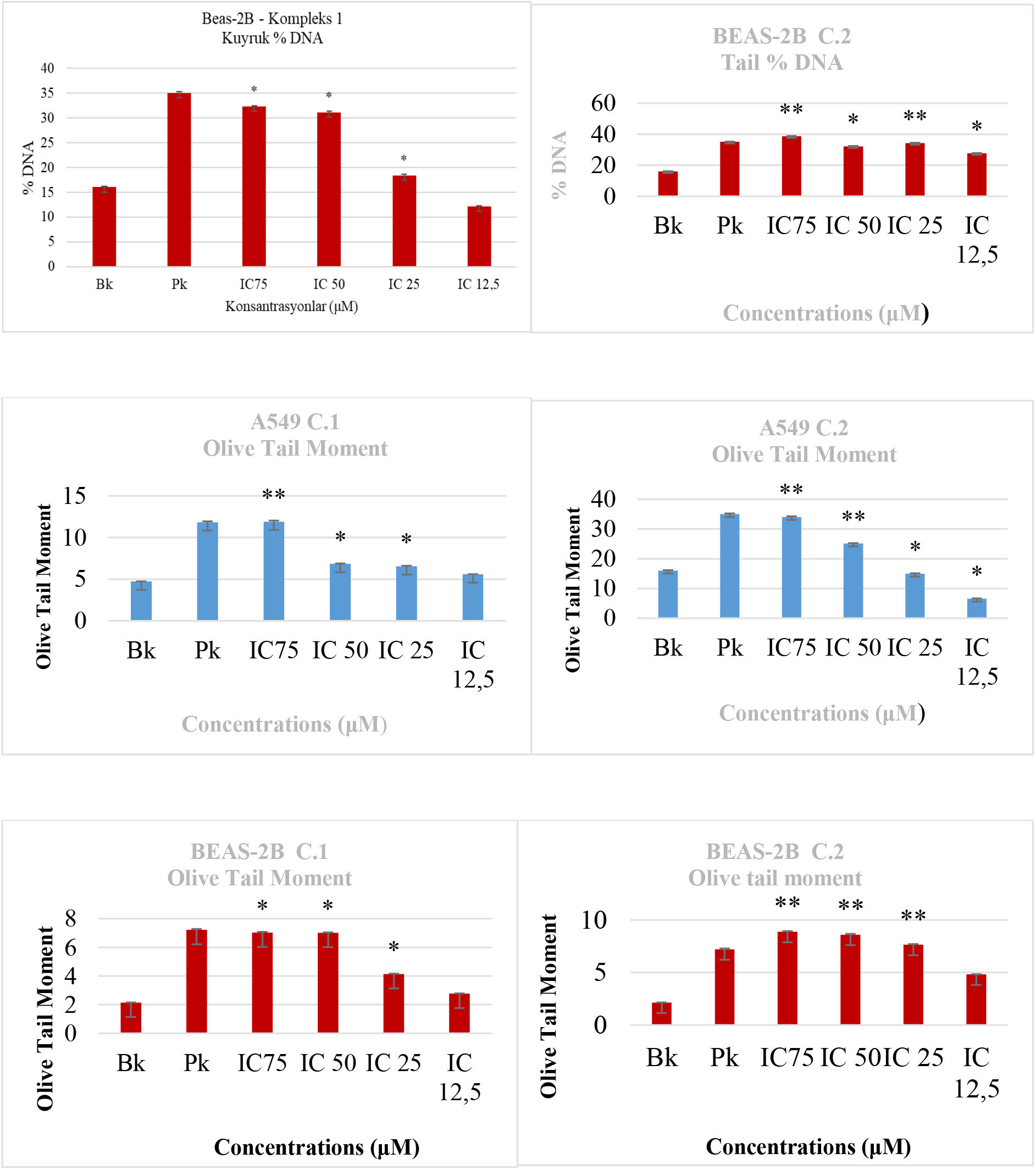
Comet assay results in A549 and BEAS-2B cell lines exposed to the complexes 1–2 (* p < 0.05, **p < 0.001)

